# A single high-zinc activation enhancer can control two genes orientated head-to-head in *C. elegans*

**DOI:** 10.1101/2024.11.19.624376

**Authors:** Hanwenheng Liu, Brian Earley, Adelita Mendoza, Patrick Hunt, Sean Teng, Daniel L. Schneider, Kerry Kornfeld

## Abstract

Enhancers play critical roles in gene expression, but a full understanding of their complex functions has yet to be defined. The cellular response to excess zinc levels in *C. elegans* requires the HIZR-1 transcription factor, which binds the high-zinc activation (HZA) enhancer in the promoters of multiple target genes. Cadmium hijacks the excess zinc response by binding and activating HIZR-1. By analyzing the genome-wide transcriptional response to excess zinc and cadmium, we identified two positions in the genome where head-to-head oriented genes are both induced by metals. In both examples, a single predicted HZA enhancer is positioned between the two translational start sites. We hypothesized that a single enhancer can control both head-to-head genes, an arrangement that has not been extensively characterized. To test this hypothesis, we used CRISPR genome editing to precisely delete the HZA^mT^ enhancer positioned between *mtl-2* and *T08G5.1*; in this mutant, both head-to-head genes display severely reduced zinc-activated transcription, whereas zinc-activated transcription of more distant genes was not strongly affected. Deleting the HZA^cF^ enhancer positioned between *cdr-1* and *F35E8.10* caused both head-to-head genes to display reduced cadmium-activated transcription, whereas cadmium-activated transcription of more distant genes was not strongly affected. These studies rigorously document that a single HZA enhancer can control two head-to-head genes, advancing our understanding of the diverse functions of enhancers.

**Article Summary:** Enhancers are critical for gene expression, but a full understanding their functions has yet to be elucidated. We discovered two positions in the *C. elegans* genome where a pair of head-to-head oriented genes are both transcriptionally activated by excess zinc and/or cadmium; in both cases, one high-zinc activation (HZA) enhancer is positioned between the translation start sites. When these HZA enhancers were deleted by genome engineering, both head-to-head genes lost metal-activated transcription. These results demonstrate that a single HZA enhancer can control two head-to-head genes, advancing the understanding of enhancer function.

## Introduction

Zinc is an essential metal. About 6% of the prokaryotic proteome and 10% of the eukaryotic proteome requires zinc for structural, catalytic, and signaling functions (Andreini et al. 2009). In humans, zinc is required for normal functioning of multiple systems, such as the nervous, immune, and reproductive systems, in addition to growth and development (Coleman 1992; Hambidge 2000; Sandstead 2015; Zoroddu et al. 2019). Both a lack and an excess of zinc lead to defects and diseases. For *C. elegans*, both deficient and excess zinc in growth media retard growth (Davis et al. 2009; Mendoza et al. 2024). In humans, zinc deficiency leads to infantile morbidity and symptoms shown in acrodermatitis enteropathica, a genetic disorder that affects dermal, digestive, immune, and reproductive systems; excess zinc is neurotoxic and is one of the causes for metal fume fever (Aggett 1989; Black 2001; Mezzaroba et al. 2019; Schoofs et al. 2024; Brenner and Keyes 2025). To prevent these adverse effects, organisms have evolved robust mechanisms to maintain appropriate cellular zinc levels. In *C. elegans*, HIZR-1, a nuclear receptor, functions as a zinc excess sensor and a transcription factor for zinc-inducible genes.

HIZR-1 senses zinc through direct binding to the ligand-binding domain; HIZR-1 activates transcription through direct binding to the HZA (**H**igh-**Z**inc **A**ctivation) element via the DNA-binding domain. The HZA enhancer is present in the promoters of multiple genes activated by zinc, including genes encoding for zinc exporters (*cdf-2*, *ttm-1*), metallothioneins (*mtl-1*, *mtl-2*), and itself (*hizr-1*) (Roh et al. 2015; Warnhoff et al. 2017). Increased expression of these proteins expels, sequesters, or chelates excess zinc to restore zinc homeostasis (Davis et al. 2009; Hall et al. 2012; Roh et al. 2013; Warnhoff et al. 2017; Essig et al. 2024).

Cadmium is a potent environmental heavy-metal toxicant. Cadmium and zinc have chemical similarities, since both are divalent d-block elements belonging to group 12 of the periodic table. Exposure to cadmium causes transcriptional changes in stress-response genes and metal chelator genes (Jin et al. 2003; Lützen et al. 2004; Cui et al. 2007; Li et al. 2015; Nordberg et al. 2015). Earley et al (2021) showed that cadmium directly binds the HIZR-1 ligand-binding domain, similar to zinc. While cadmium has been proposed to displace physiological metals and cause protein dysfunction, in the case of HIZR-1, cadmium promotes nuclear accumulation in intestinal cells and activation of HIZR-1-dependent transcription via the HZA enhancer. The dramatic transcriptional response to cadmium exposure that is mediated by HIZR-1 indicates that cadmium can function as a zinc mimetic to activate the high zinc homeostasis pathway.

Enhancers are *cis-*regulatory DNA sequences that either enhance or repress gene expression (Banerji et al. 1981; Levine 2010). By recruiting various transcription factors, enhancers drive cell differentiation and regulate stress response pathways (Panigrahi and O’Malley 2021). Enhancers function at various locations relative to their target genes. They can be located either up- or downstream of the target genes, as close as less than 1kb or as far as more than 100kb away. A single enhancer may regulate multiple genes (Chen et al. 2013; Mills et al. 2020). Usually, these genes are encoded on the same DNA strand in a series, and the enhancer is positioned either 5’ or 3’ of the series of genes, similar to operons in bacteria (Lercher et al. 2003; Cutter et al. 2009). When two genes that are encoded on different DNA strands flank a common 5’ region, they are called head-to-head genes. Head-to-head genes are widespread in the *C. elegans* and human genomes; in these cases, transcription of both genes is initiated from the common 5’ region. This is called “divergent transcription” (Adachi and Lieber 2002; Trinklein et al. 2004; Seila et al. 2009; Ibrahim et al. 2018). The common 5’ regions may contain enhancers that in principle could regulate one or both transcripts. However, few examples of this situation have been analyzed in detail. Examples of head-to-head gene pairs that have been analyzed in mice and chordates indicate that enhancers in the common 5’ region regulate one gene or the other, but not both, which ensures cell type-specific gene expression (Swamynathan and Piatigorsky 2002; Hozumi et al. 2013). Whether this logic also applies to the HZA enhancer in *C. elegans* is unknown.

The function of the HZA enhancer was previously analyzed using transgenic animals with extrachromosomal arrays — in this context, the HZA enhancer was demonstrated to be necessary for the zinc-induced activation of the *mtl-2* gene and sufficient to confer zinc-induced activation on a *pes-10* basal promoter (Roh et al. 2015). However, the function of endogenous HZA enhancers has not been reported. Here we used CRISPR genome editing to directly evaluate the function of endogenous HZA enhancers. We examined two examples of head-to-head genes in *C. elegans* that are both activated by cadmium (Earley et al. 2021). Our results indicate that a single HZA enhancer can control metal-activated transcription of both head-to-head oriented genes. Thus, the regulatory function of a single HZA enhancer is divergent and acts on two direct target genes, elucidating the functional capacity of this important DNA control element.

## Methods and materials

### Worm Handling and Culture

Animals were cultured on nematode growth media (NGM) dishes seeded with *E. coli* OP50 at 20℃ (Brenner 1974) unless otherwise noted. We picked three L1-stage larvae onto fresh NGM dishes weekly to maintain strains.

To minimize precipitation of metals during supplementation experiments, we cultured animals on noble agar minimal media (NAMM) dishes (Warnhoff et al. 2017; Earley et al. 2021). Each NAMM dish contains 0.1% cholesterol and 3.7% noble agar. Zinc replete NAMM has no supplemental metal. Zinc excess NAMM contains 200μM ZnSO_4_. Cadmium NAMM contains 100μM CdCl_2_.

### Strain generation

Strains used in this study are listed in Table S1. We used Bristol isolate N2 as WT (Brenner 1974). The *hizr-1(am286)* strain, WU1958, was generated by outcrossing the parental strain, WU1500, to N2 three times. The *am286* allele contains a C259T mutation that changes Q87 to STOP (Warnhoff et al. 2017).

We worked with SunyBiotech (www.sunybiotech.com) to generate CRISPR knockout strains, PHX4134 and PHX4265. PHX4134 contains *syb4134*, a 14bp deletion that removes 14bp of the HZA^cF^ element (AAC AGA AAC TAC AA) that is positioned 108bp upstream of the *cdr-1* translation start site (ATG) (Fig. 3). PHX4265 contains *syb4265*, a 15bp deletion that removes the entire 15bp HZA^mT^ element (ATC ACA AAC TAG AGT) that is positioned 278bp upstream of the *mtl-2* translation start site (ATG) (Fig. 2). We validated the position of these deletions by Sanger DNA sequencing.

To analyze the *mtl-2(gk125)* mutation, we outcrossed VC128 five times to N2 to obtain the strain WU946 (The C. elegans Deletion Mutant Consortium 2012). The *gk125* insertion/deletion removes the region from 208bp upstream of *mtl-2* ATG to 584bp downstream of *mtl-2* STOP codon (TAA) and inserts a single adenine (Fig. 4). We validated the position of the insertion/deletion by Sanger DNA sequencing. DNA sequencing was performed using GENEWIZ (Azenta Life Sciences, www.genewiz.com). See Table S2 for oligonucleotide primers.

### RNA extraction, qPCR, and statistical analysis

To age-synchronize the animals, we grew each strain on 90mm NGM dishes until the population was recently starved and most animals were adult stage. We washed the animals off the NGM dishes with M9 buffer into a 14ml falcon tube (M9 buffer: 1ml 1M MgSO_4_, 3g KH_2_PO_4_, 6g Na_2_HPO_4_, 5g NaCl in 1L MilliQ (Stiernagle 2006)). We centrifuged and washed the worms with M9 buffer three times before one final centrifugation for a pellet. We added 5ml bleaching solution to the pellet and vortexed for three minutes to release the eggs (bleaching solution: MilliQ, commercial Clorox bleach, and 1M NaOH at a ratio of 7:2:1). If live worms were still present, we extended vortexing for another 15 seconds. We washed the eggs with M9 buffer twice and hatched them on 90mm unseeded NGM dishes. After 24 hours, we transferred the hatched L1 larvae onto 90mm seeded NGM dishes and cultured for 28 to 32 hours until most animals were L4 stage. To avoid crowding, which may reduce the growth rate, we sometimes used two dishes. L4 animals were separated onto NAMM dishes corresponding to different metal conditions. After 16 hours, we collected the animals for RNA extraction according to previous publications (Warnhoff et al. 2017; Earley et al. 2021). Briefly, we washed animals from NAMM dishes to an Eppendorf tube, pelleted to 100μl, and added 400μl TRIzol (Invitrogen Cat# 15596018). We subjected the tube to a liquid nitrogen-37℃ heatblock freeze cycle three times, added an additional 200μl TRIzol and 140μl chloroform (Sigma-Aldrich Cat#319988), vortexed, and centrifuged. We purified the top aqueous phase containing crude RNA extract using Qiagen RNeasy Mini Kit (Cat# 74106). For cDNA synthesis, we used 1μg RNA and followed instruction from Intact Genomics igScript First Strand cDNA Synthesis Kit (Cat# 4314) or BioRad iScript reverse transcription supermix for RT-qPCR (Cat# 1708841). We substituted the water with Ambion nuclease-free water (Invitrogen Cat# AM9937). We performed qPCR using MicroAmp fast optical 96-well reaction plate (ThermoFisher Cat# 4346906) and BioRad iTaq Universal SYBR Green Supermix (Cat# 1725121) in an Applied Biosystems StepOne Plus thermocycler. The thermocycling program was 95℃ for 20 seconds, then 40 cycles of 95℃ for 3 seconds, and 60℃ for 30 seconds. Primers for qPCR were designed with the IDT PrimerQuest Tool (www.idtdna.com/PrimerQuest). The primers are listed in Table S2. For each biological replicate, we measured three technical replicates for each gene (three wells). The results were collected as C_T_ cycles and analyzed with 2^-ΔΔC^_T_ method in Microsoft Excel (Schmittgen and Livak 2008).

GraphPad Prism 9 was used for statistical analysis and graphing. We first transformed the 2^-ΔΔC^_T_ values of each technical replicate by base 2 logarithm for normal distribution, and then removed outliers with the ROUT method (Q=0.5%). We calculated the biological replicate data points as averages of the respective outlier-removed technical replicates. We analyzed the biological data with ANOVA or mixed-effects analysis with Holm-Šídák correction. Changes in expression levels of any given gene are reported as the average of all biological data in fold change; “3 fold” means 2^3^, or “8 times”. We overlaid both technical and biological data into a single SuperPlot in Adobe Illustrator (Lord et al. 2020).

## Results

### Identification of two genomic regions containing adjacent genes that are oriented head-to-head and regulated by cadmium

Earley et al. (2021) identified many *C. elegans* genes that are activated by cadmium in wild-type (WT) animals. While analyzing the chromosomal locations of these genes, we noticed that a region of chromosome V (14,018,240-14,020,360) contains two activated genes that are oriented head-to-head. One gene is *mtl-2,* which encodes a *C. elegans* metallothionein protein that has been characterized extensively for its roles in cadmium and zinc biology (Bofill et al. 2009; Zeitoun-Ghandour et al. 2010; Hall et al. 2012; Essig et al. 2024). The other gene is *T08G5.1,* which encodes an uncharacterized protein. Roh et al. (2015) reported that the *mtl-2* promoter contains a predicted HZA enhancer that promotes gene activation by zinc and cadmium. This HZA element, named “HZA^mT^” hereafter, is positioned between these two genes: the start codon ATGs of *mtl-2* and *T08G5.1* are 277bp and 288bp from the HZA^mT^ element, respectively (Fig. 1A). To quantify the transcriptional response to cadmium, we exposed WT animals at the L4 stage to 100μM cadmium for 16 hours, prepared RNA, and determined mRNA levels by quantitative reverse transcriptase polymerase chain reaction (qRT-PCR) (Earley et al. 2021). Compared to control worms not exposed to cadmium, *mtl-2* and *T08G5.1* mRNA levels were significantly increased by 5.4 log2 fold (∼42x) and 7.6 log2 fold (∼190x), respectively (Fig. 1B-D).

**Figure 1.**
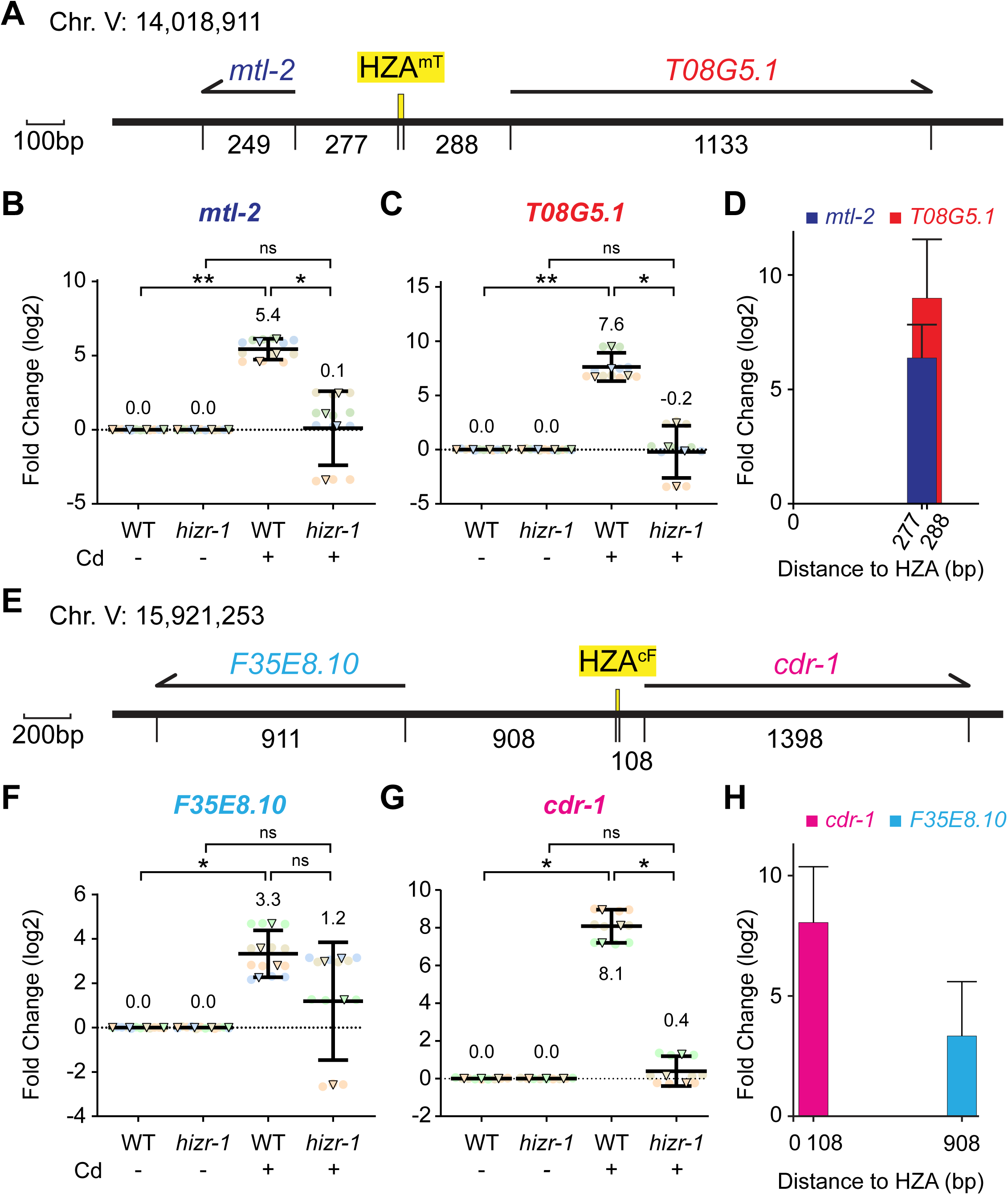
Two regions of Chromosome V have genes that are oriented head-to-head, activated by cadmium, and require *hizr-1* for activation. **A, E)** Schematics of regions of Chromosome V are drawn to scale in base pairs (bp). Numeric chromosome location indicates the first base pair of the HZA. Thick black line represents DNA; yellow box indicates HZA enhancer; numbers are intervals from HZA enhancer to ATG start codon and end of transcript; arrows above represent pre-mRNA for indicated genes. Scale bar as shown. **B, C, F, G)** Wild type (WT) and *hizr-1(am286)* mutants at the L4 stage were cultured with or without 100μM cadmium for 16 hours and analyzed by qPCR. Values for WT and *hizr-1(am286)* with no cadmium were set equal to 0, and values with cadmium represent log2 fold change. N = 4 initial biological replicates but may vary in panels due to outlier removal. Circles are technical replicates (repeated measurements of the same sample), and triangles are biological replicates (different samples from different days). Same color denotes the same experiment trial. Statistical analysis by pairwise one-way ANOVA. Non-significant p-values are listed for p<0.3; otherwise, “ns”. For significant p-values: *<0.05; **<0.01; ***<0.001; ****<0.0001. Error bars represent mean ± 95% confidence intervals. Mean values are listed. All fold changes are in base 2 logarithm (e.g. 3 folds = 8x). The same approach is used to represent qPCR data in Figures 2-4, S1-S3. **D, H)** Distances from the HZA to the start codons (shown in panels A and E) plotted against fold change for WT plus cadmium (shown in B, C, F, and G).

We noticed another example of two genes that are oriented head-to-head and both activated by cadmium on chromosome V (15,919,280-15,922,860). One gene is *cdr-1*, which was identified and named after its strong induction by cadmium (Hall et al. 2012). The other gene is *F35E8.10*, which encodes an uncharacterized protein. By searching the region between these two genes, Roh et al. (2015) identified a predicted HZA element, named “HZA^cF^” hereafter: the start codon ATGs of *F35E8.10* and *cdr-1* are 908bp and 108bp from the HZA^cF^ element, respectively (Fig. 1E). Compared to control worms not exposed to cadmium, *F35E8.10* and *cdr-1* mRNA levels were significantly increased by 3.3 log2 fold (∼10x) and 8.1 log2 fold (∼270x), respectively (Fig. 1F-G). For these two genes, the increase in mRNA levels correlated with the distance from the predicted HZA element (Fig. 1H).

### *hizr-1* was necessary for cadmium-mediated transcript accumulation of *mtl-2*, *T08G5.1, cdr-1,* and *F35E8.10*

Warnhoff et al. (2017) showed that the HZA-binding transcription factor, HIZR-1, is required for zinc-induced expression of multiple genes. To test the model that HIZR-1 is necessary for activation of these four head-to-head genes in response to cadmium, we used qPCR to quantify transcript levels in a strong loss-of-function mutant, *hizr-1(am286* Q87STOP) (Warnhoff et al. 2017). Indeed, all four genes were affected. *mtl-2*, *T08G5.1, F35E8.10,* and *cdr-1* mRNA levels were not significantly different in *hizr-1* mutants treated with cadmium or untreated. In addition, transcript levels of *mtl-2*, *T08G5.1,* and *cdr-1* were significantly lower in *hizr-1* mutants treated with cadmium compared to WT animals treated with cadmium (Fig. 1).

*F35E8.10* transcript levels were lower in *hizr-1* mutants treated with cadmium compared to WT animals treated with cadmium, but this trend was not significant with this sample size (Fig. 1F). Thus, there may be a *hizr-1*-independent mechanism to activate transcription of *F35E8.10* in response to cadmium. These results are consistent with previous reports (Roh et al. 2015; Earley et al. 2021) and indicate *hizr-1* is necessary, at least in part, for the cadmium-induced accumulation of the transcripts these genes.

Earley et al (2021) examined the role of *hizr-1* in the response of three of these genes to excess zinc: *mtl-2*, *T08G5.1,* and *cdr-1.* RNA-seq analysis showed that mRNA accumulation of these three genes is significantly increased by excess zinc, and this response is strongly reduced or abrogated in *hizr-1(lf)* mutants (Earley et al. 2021). The function of *hizr-1* in regulating the level of *F35E8.10* mRNA in response to excess zinc has not been reported.

### A single HZA element mediates transcriptional activation of *mtl-2* and *T08G5.1* in response to zinc

Because HIZR-1 directly binds the HZA element, we hypothesized that the HZA elements mediated transcript accumulation in response to metal. To evaluate this model, we used CRISPR/Cas9 to generate the *syb4265* mutation, a 15-bp deletion that removes the HZA^mT^ element on chromosome V at position 14,018,911 (Fig. 2B; labeled “HZA^mT^Δ”). The sequence and the approximate position of this HZA enhancer in the *mtl-2* promoter have been conserved in the *Caenorhabditis* genus (Fig. 2B). Because zinc is the physiological metal that activates HIZR-1, we first examined the response to excess zinc. WT and HZA^mT^Δ mutant animals at the L4 stage were cultured on noble agar minimal media (NAMM) with either no additional metals (replete zinc) or 200μM supplemental zinc (excess zinc) for 16 hours, and gene expression levels were quantified by qPCR. Compared to control worms in zinc replete medium, WT animals in zinc excess medium displayed *mtl-2* and *T08G5.1* mRNA levels that were significantly increased by 3.7 log2 fold (∼13x) and 4.2 log2 fold (∼18x), respectively (Fig. 2C-D). By contrast, *mtl-2* and *T08G5.1* mRNA levels were not significantly different in HZA^mT^Δ mutants cultured in zinc replete or excess medium. In addition, transcript levels were significantly lower in HZA^mT^Δ mutants treated with excess zinc compared to WT animals treated with excess zinc (Fig. 2C-D). Thus, the HZA^mT^ enhancer was necessary for zinc-activated transcription of both genes.

**Figure 2.**
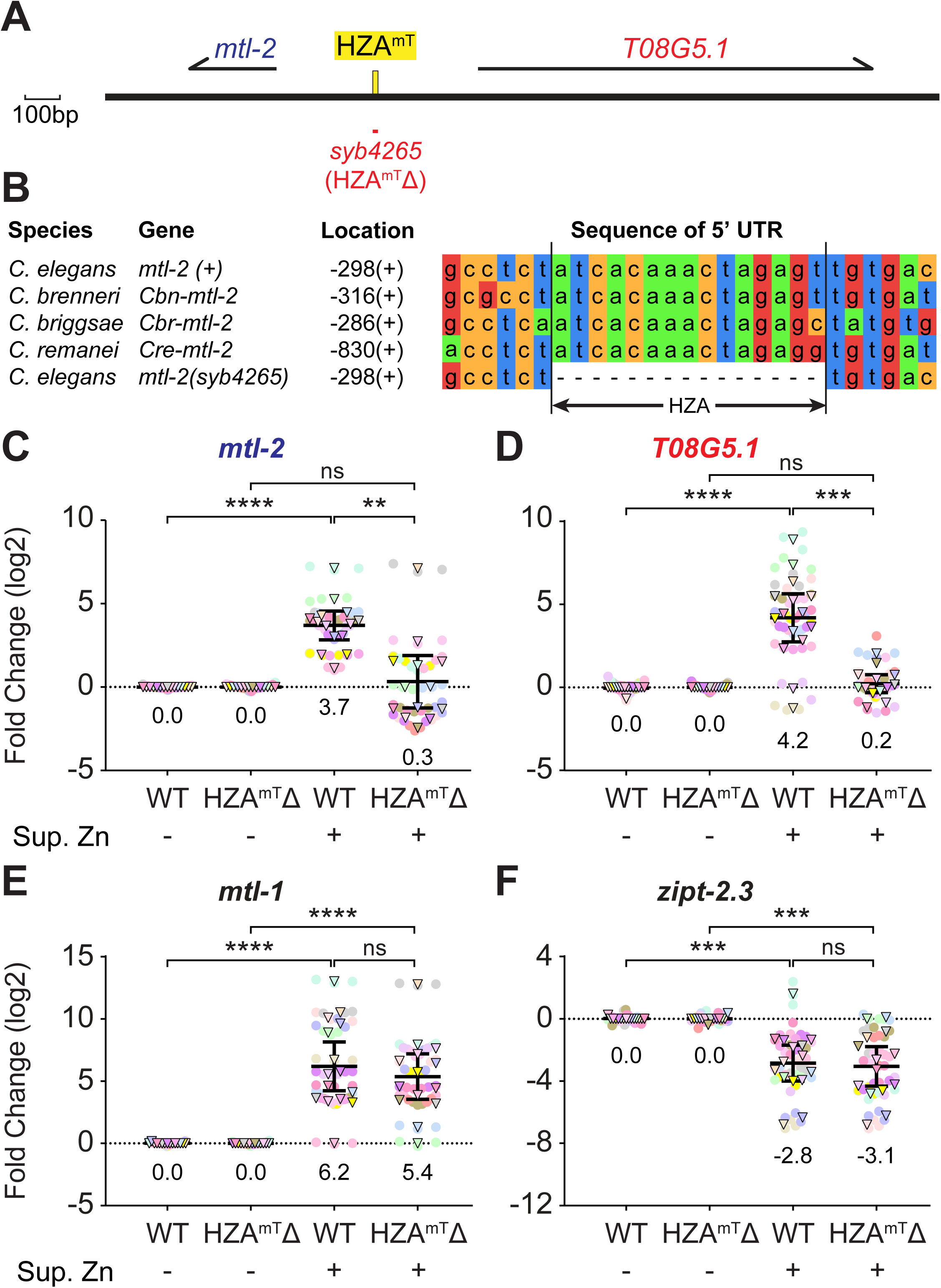
The HZA^mT^ element is necessary for zinc-induced transcription of both *mtl-2* and *T08G5.1*. **A)** Schematic of a region of Chromosome V drawn to scale in base pairs (bp). Thick black line represents DNA; yellow box indicates HZA enhancer. Arrows above represent pre-mRNA for indicated genes. Red line below indicates mutation *syb4265*, a 15-bp deletion of the HZA^mT^ (named HZA^mT^Δ). Scale bar: 100bp. **B)** Alignment of DNA sequences from the 5’ untranslated regions of *mtl-2* genes, containing the conserved HZA^mT^ element and six flanking base pairs. Sequences are from four WT *Caenorhabditis* species and the *C. elegans syb4265* deletion strain. Location indicates the number of base pairs upstream from the ATG start codon and the strand orientation. **C-E)** Wild type (WT) and *syb4265* mutants at the L4 stage were cultured with replete (-) or 200μM supplemental zinc (+) for 16 hours and analyzed by qPCR. N = 15 initial biological replicates but may vary in panels due to outlier removal.

To determine the specificity of the transcriptional response to the deleted HZA^mT^ enhancer, we examined zinc-regulated genes at distant genomic positions (Fig. 2E-F, S1). Similar to WT animals in excess zinc, HZA^mT^Δ mutants in excess zinc displayed significantly higher transcript levels for two activated genes on chromosome X: *cdf-2* (chrX:12,462,640-12,468,917) and *hizr-1* (chrX:11,316,278-11,323,086); and two activated genes on chromosome V: *mtl-1* (chrV:6,689,370-6,693,863) and *cdr-1* (chrV: 15,921,309-15,922,820) (Fig. S1). Furthermore, *zipt-2.3* (chrII:13,710,905-13,717,893) on chromosome II that is repressed by excess zinc was not significantly affected by this mutation (Fig. 2F). These results indicate that the HZA^mT^Δ mutation specifically affected the response to excess zinc in adjacent genes, but not in genes positioned distantly on the same chromosome or genes on different chromosomes. Thus, neither the HZA^mT^ enhancer itself, nor the activation of *mtl-2* and *T08G5.1* by excess zinc mediated by the HZA^mT^ enhancer, was necessary for the robust regulation by excess zinc of five distantly positioned genes.

To evaluate the function of the HZA^mT^ enhancer in response to cadmium, we exposed WT and HZA^mT^Δ mutant animals at the L4 stage to NAMM with 0 or 100μM cadmium for 16 hours and determined mRNA levels by qPCR. In WT animals, both *mtl-2* and *T08G5.1* mRNA levels were significantly increased by cadmium exposure (Fig. S2A-B). By contrast, HZA^mT^Δ mutant animals displayed much less robust levels of *mtl-2* and *T08G5.1* activation that were not significant. Thus, the HZA^mT^ enhancer was necessary for full cadmium-activated transcription of both flanking genes. However, the residual activation of these two genes by cadmium suggests there may be an activation mechanism that is independent of the HZA^mT^ enhancer.

Unexpectedly, we observed a partial reduction of activation with four control genes positioned far from the mutation: *cdf-2*, *mtl-1*, *cdr-1,* and *F35E8.10* (Fig. S2C-F). These results indicate that the HZA^mT^Δ mutation affected the response to cadmium in adjacent genes, genes positioned distantly on the same chromosome, and genes on different chromosomes. Thus, either the HZA^mT^ enhancer itself, or cadmium activation of *mtl-2* and *T08G5.1* mediated by the HZA^mT^ enhancer, was necessary for the robust regulation by cadmium of four distantly positioned genes. Another possible interpretation is that a background mutation in the HZA^mT^Δ strain reduces the robustness of cadmium-activated transcription throughout the genome.

### A single HZA element mediates transcriptional activation of *cdr-1* and *F35E8.10* in response to cadmium

We used CRISPR/Cas9 to delete the HZA element between *cdr-1* and *F35E8.10*; the *syb4134* mutation is a 14-bp deletion that removes 14bp of the HZA^cF^ element (Fig. 3A-B; labeled as “HZA^cF^Δ”). To evaluate the response of these genes to zinc, we quantified gene expression levels of the WT and HZA^cF^Δ animals as described above. Compared to WT animals on zinc replete medium, WT animals in zinc excess displayed *cdr-1* mRNA levels that were significantly increased by 7.5 log2 fold (∼180x) (Fig. S3B). In HZA^cF^Δ mutants, *cdr-1* mRNA levels were not significantly different when treated with excess zinc or untreated. In addition, *cdr-1* transcript levels were significantly lower in HZA^cF^Δ mutants treated with excess zinc compared to WT animals treated with excess zinc (Fig. S3B). Thus, the HZA^cF^ element was necessary for zinc-induced transcription of *cdr-1*. Compared to WT animals on zinc replete medium, WT animals in zinc excess displayed *F35E8.10* mRNA levels increased by 1.7 log2 fold (3.2x), but this trend was not significant with this sample size (Fig. S3A). In HZA^cF^Δ mutants, *F35E8.10* mRNA levels were not significantly different when treated with excess zinc or untreated. Because excess zinc did not significantly activate *F35E8.10* in WT, we cannot use the analysis of the HZA^cF^Δ mutant to determine if the enhancer is necessary.

**Figure 3.**
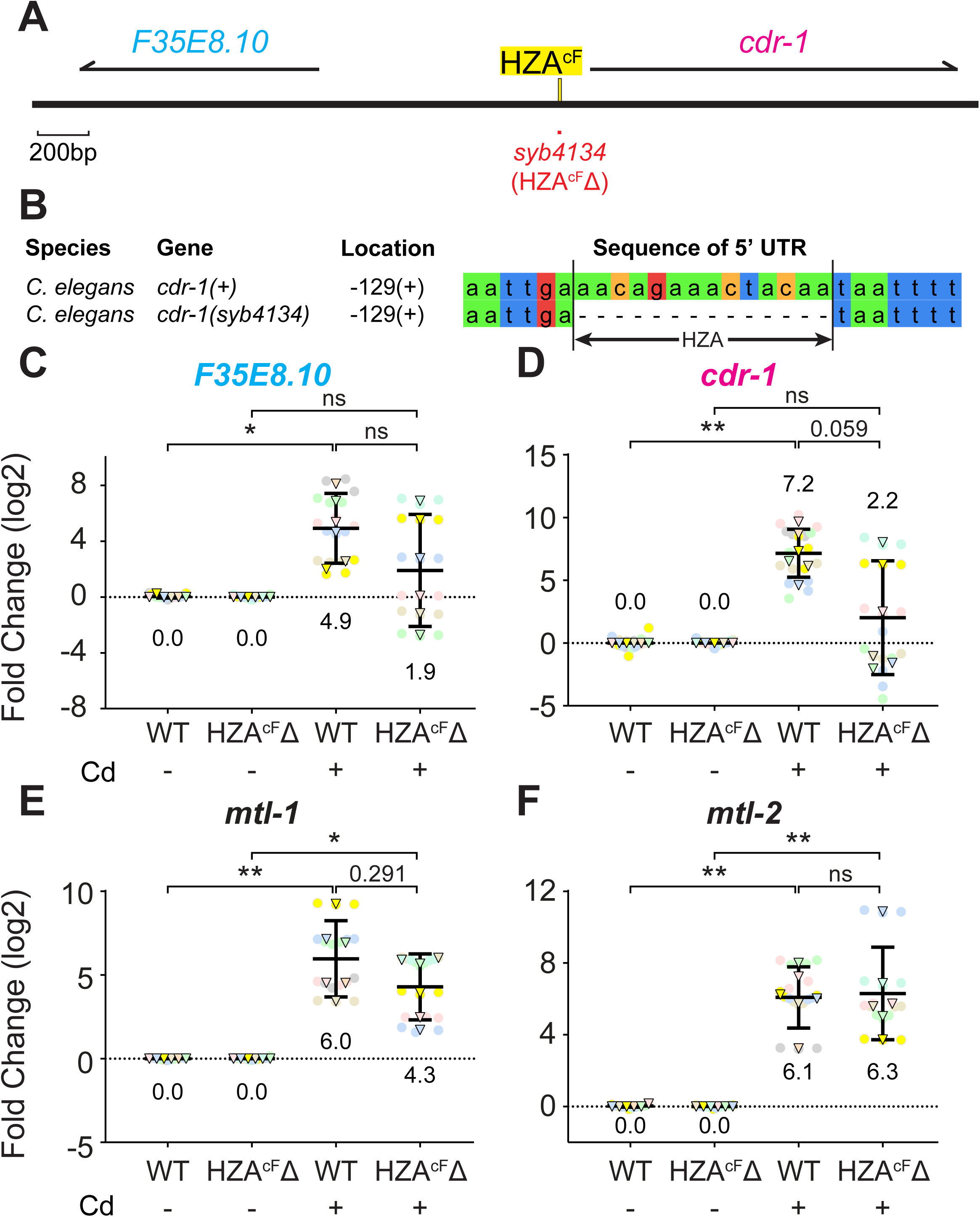
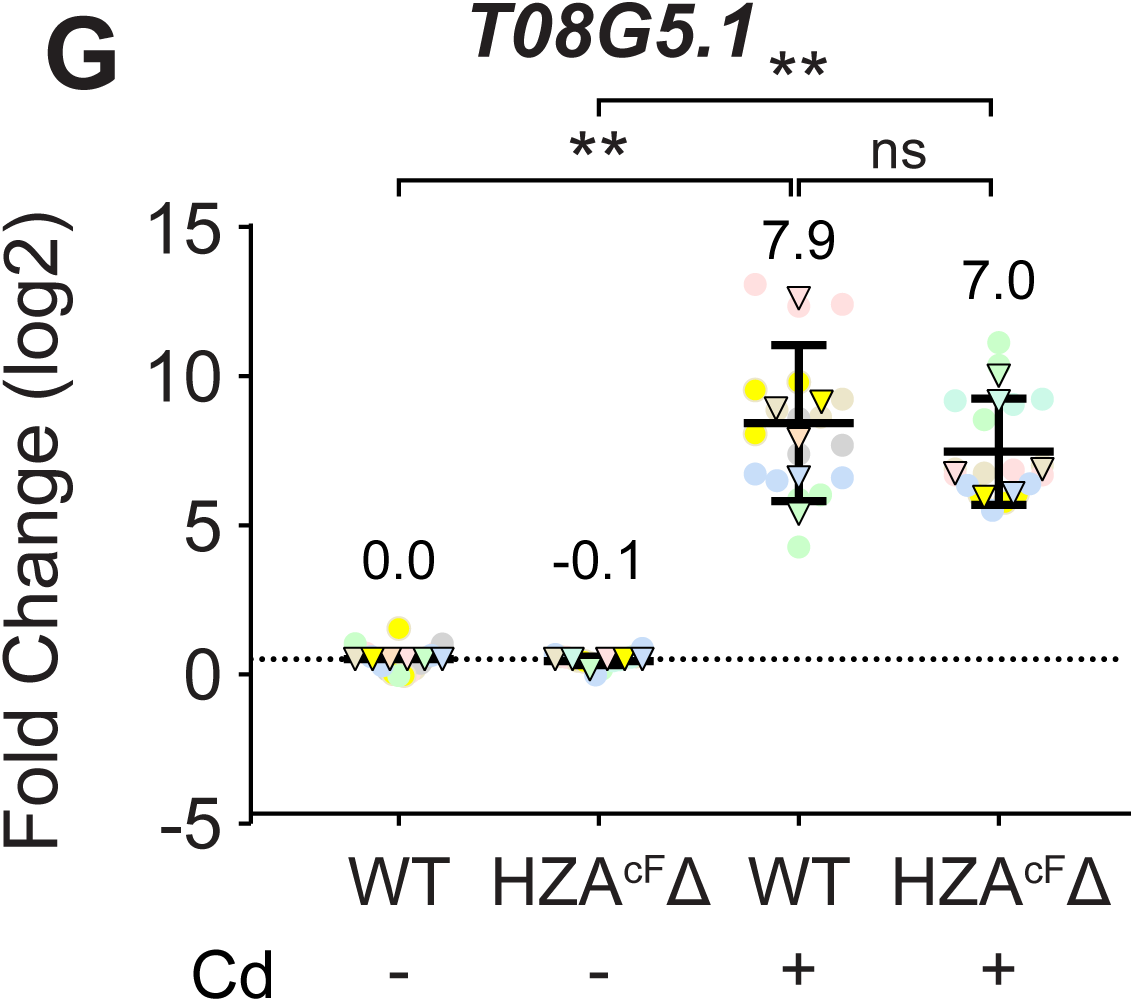
The HZA^cF^ element was necessary for full cadmium-induced transcript accumulation of both *cdr-1* and *F35E8.10*. **A)** Schematic of a region of Chromosome V drawn to scale in base pairs (bp). Thick black line represents DNA; yellow box indicates HZA enhancer. Arrows above represent pre-mRNA for indicated genes. Red line below indicates the *syb4134* mutation, a 14-bp deletion of the HZA^cF^ enhancer (named HZA^cF^Δ). Scale bar: 200bp. **B)** Alignment of DNA sequences from the 5’ untranslated regions of *cdr-1* that contain the conserved HZA^cF^ element and six flanking base pairs in WT and *syb4134* mutant. Location indicates the number of base pairs upstream from the ATG start codon and the strand orientation. **C-G)** Wild type (WT) and *syb4134* mutants at the L4 stage were cultured with (+) or without (-) 100μM cadmium for 16 hours and analyzed by qPCR. N = 6 initial biological replicates, but may vary in panels due to outlier removal.

To determine the specificity of the transcriptional response to the deleted HZA^cF^ enhancer, we examined control genes at distant genomic positions (Fig. S3C-H). Similar to WT animals in excess zinc, HZA^cF^Δ mutants in excess zinc displayed significantly higher transcript levels for two activated genes on chromosome X: *cdf-2* and *hizr-1*; and three activated genes on chromosome V: *mtl-1*, *mtl-2* (chrV: 14,018,270-14,018,673), and *T08G5.1* (chrV:14,018,766-14,020,388). Furthermore, *zipt-2.3* on chromosome II that is repressed by excess zinc was not significantly affected by this mutation. These results indicate that the HZA^cF^Δ mutation specifically affected the response to excess zinc in the adjacent gene *cdr-1*, whereas it did not affect genes positioned distantly on the same chromosome or genes on different chromosomes. Thus, neither the HZA^cF^ enhancer itself, nor the activation of *cdr-1* by excess zinc mediated by the HZA^cF^ enhancer, was necessary for the robust regulation by excess zinc of six distantly positioned genes.

To evaluate the effect of the deleted HZA^cF^ enhancer on the response to cadmium, we analyzed WT and HZA^cF^Δ mutant animals as described above. Compared to control worms cultured without cadmium, WT animals cultured with cadmium displayed *F35E8.10* and *cdr-1* mRNA levels that were significantly increased by 4.9 log2 fold (∼30x) and 7.2 log2 fold (∼147x), respectively (Fig. 3C-D). By contrast, in HZA^cF^Δ mutants *F35E8.10* and *cdr-1* did not display significant induction, indicating the HZA^cF^ element is necessary for full transcriptional regulation. However, *cdr-1* and *F35E8.10* did display a trend towards weak induction that was not significant with this sample size, and the level of induction was not significantly different in WT and HZA^cF^Δ mutants. These results suggest the existence of an HZA^cF^-independent mechanism for cadmium-mediated induction of *cdr-1* and *F35E8.10.* For control genes positioned far from the mutation on chromosome V, *mtl-1*, *mtl-2*, and *T08G5.1,* transcriptional activation by cadmium was similar in WT and HZA^cF^Δ mutant animals (Fig. 3E-G). Thus, deleting the HZA^cF^ enhancer significantly reduced cadmium activation of both adjacent genes, whereas it minimally affected cadmium activation of distant genes on the same chromosome.

### A deletion of *mtl-2* did not influence transcription of *T08G5.1*

Our results establish that *mtl-2* and *T08G5.1* are regulated by the same HZA enhancer. If these two genes compete for access to the HZA^mT^ element, then removing one gene might increase transcription of the other. No deletions of *T08G5.1* currently exist, but we were able to take advantage of the existing mutation, *mtl-2(gk125*), which deletes a region from 208bp upstream of the *mtl-2* translation start codon (ATG) to 584bp downstream of the *mtl-2* STOP codon and inserts one adenine (Fig. 4A-B; labeled “*mtl-2Δ*”) (Hall et al. 2012; The C. elegans Deletion Mutant Consortium 2012). The deleted region includes the predicted TATA box (TATAAAAG) and a GATA element (CTGATAA), positioned 47bp and 87bp upstream of *mtl-2* ATG, respectively (Fig. 4B). GATA elements are tissue-specific enhancers that promote expression in the intestine and function with the HZA to drive the transcriptional response to excess zinc in intestinal cells (Moilanen et al. 1999; Roh et al. 2015). The HZA^mT^ element and a second GATA element are not deleted. As expected, *mtl-2* mRNA was not reliably detected in *mtl-2Δ* mutants. We evaluated the response of WT and *mtl-2Δ* mutants to excess zinc as described above. Compared to animals in zinc-replete medium, WT and *mtl-2Δ* mutants in zinc excess medium displayed *T08G5.1* mRNA levels that were significantly increased by 4.6 log2 fold (∼24x) and 4.7 log2 fold (∼26x) (Fig. 4C). Thus, deletion of *mtl-2* did not affect transcriptional activation of *T08G5.1* in response to zinc excess.

**Figure 4.**
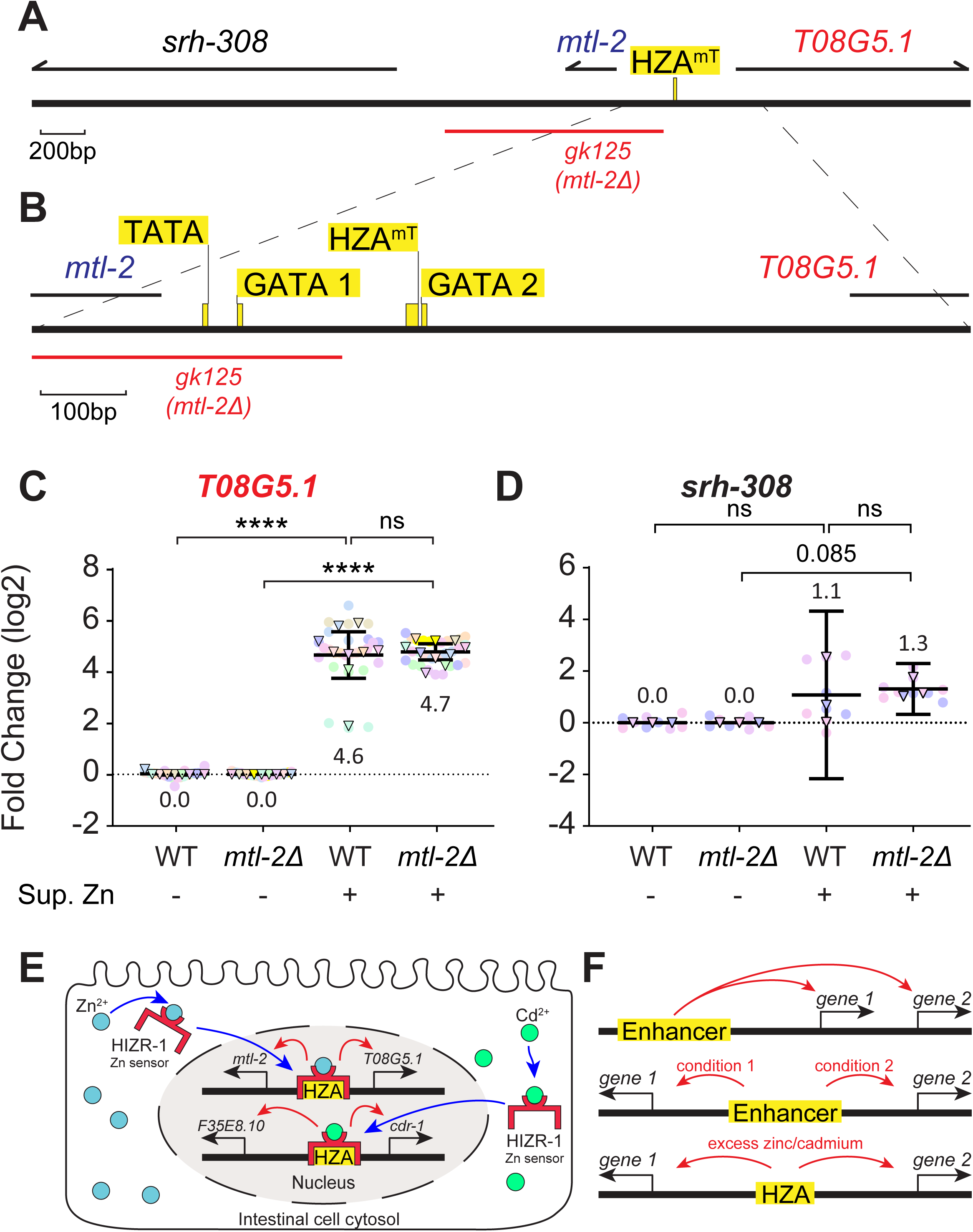
A deletion of *mtl-2* did not influence zinc-activated transcription of *T08G5.1*. **A)** Schematic of a region of Chromosome V drawn to scale in base pairs (bp). Thick black line represents DNA; yellow box indicates HZA enhancer. Arrows above represent pre-mRNA for indicated genes. Red line below indicates the extent of the *gk125* deletion that removes the *mtl-2* open reading frame (labeled *mtl-2Δ*). Scale bar: 200bp. **B)** An enlarged region of panel A is indicated by dashed lines and shows the positions of predicted control elements: TATA box, GATA element 1, HZA^mT^, and GATA element 2. The *gk125* deletion removes the TATA box and GATA element 1. Scale bar: 100bp. **C, D)** Wild type (WT) and *mtl-2(gk125)* mutants at the L4 stage were cultured with or without 200μM zinc for 16 hours and analyzed by qPCR. Biological replicates: *T08G5.1* N = 10, *srh-308* N = 3. **E)** Model for the orientation-independent enhancer function of the HZA element. In intestinal cells, the ligand-binding domain of HIZR-1 (red) binds to zinc ions (cyan circles) during zinc excess conditions or cadmium ions (green circles) during cadmium stress (blue arrows). HIZR-1 then enters the nucleus (blue arrows), the DNA-binding domain binds the HZA element (yellow), and transcription of both flanking genes is activated (red arrows). Thick black lines in the nucleus represent chromosome V, and thin black arrows indicate transcription start sites of genes. Not drawn to scale. **F)** Different modes of enhancer function. Top, a single enhancer controls gene 1 and gene 2, which are encoded on the same DNA strand. Middle, a single enhancer controls gene 1 in condition 1 or gene 2 in condition 2, and these head-to-head genes are encoded on different DNA strands. Bottom, a single HZA enhancer controls gene 1 and gene 2 in the same condition, and these head-to-head genes are encoded on different DNA strands.

The *mtl-2Δ* deletion results in the ATG start codon of the neighboring gene *srh-308* becoming 281bp away from the HZA^mT^ element (Fig. 4A). To evaluate the possibility that the HZA^mT^ enhancer now regulates the new adjacent gene, we examined transcription of *srh-308* as described above. *srh-308* displayed high C_T_ cycles in WT, indicating a low baseline level of transcripts (Fig. S4C). *srh-308* was not significantly induced by excess zinc in WT or *mtl-2Δ* mutants (Fig. 4D). Thus, *srh-308* does not appear to be regulated by the HZA^mT^ element in this mutant. One possible explanation for this observation is that the transcription start site of *srh-308* is deleted in *mtl-2Δ* mutants (Fig. S5) (Gu et al. 2012; Chen et al. 2013; Kruesi et al. 2013; Saito et al. 2013).

## Discussion

Based on the analysis of cadmium activated genes reported by Earley et al. (2021), we identified two examples of adjacent genes that are oriented head-to-head and both activated by cadmium. In both examples, a predicted HZA enhancer is positioned between the two genes. To address fundamental questions about the function of these HZA enhancers, we used CRISPR genome engineering to create small deletions that remove these enhancers from the genome. While genome editing technology has been used to study *cis*-regulatory elements in many organisms, including enhancers in *C. elegans*, mice, and mammalian cells, it has not previously been used to analyze the HZA enhancer (Moorthy and Mitchell 2016; Li et al. 2020; Froehlich et al. 2021).

Deleting the HZA^mT^ enhancer strongly reduced activation of both adjacent genes *mtl-2* and *T08G5.1* in response to excess zinc. By contrast, control genes positioned at a distance from this deletion that are regulated by excess zinc were minimally affected. These results lead to several conclusions: (1) within a reasonable distance, a single HZA enhancer can regulate two genes oriented head-to-head; (2) by extension, this enhancer can function in either orientation (Fig. 4E-F). Orientation independence is considered a hallmark of enhancers (Banerji et al. 1981). It is well-established biochemically that HIZR-1 binds the HZA and activates transcription (Warnhoff et al. 2017; Earley et al. 2021). However, our data do not exclude the model that another factor also binds the HZA and activates transcription. In addition to removing the HZA^mT^ enhancer, this 15 bp deletion mutation also changed the spacing, and potentially the topology, of sequence elements in the promoter (Kouzine et al. 2014). Thus, an alternative explanation for these results is that the deletion reduced zinc activated transcription as a result of this topology change. We do not favor this interpretation, because the HIZR-1 transcription factor that binds HZA enhancers to stimulate transcription is necessary for transcriptional activation of these two genes in response to excess zinc, indicating the HZA is important (Warnhoff et al. 2017; Earley et al. 2021). Furthermore, a large deletion of *mtl-2* that would change spacing and topology in the region but preserves the HZA enhancer did not affect induction of *T08G5.1* by excess zinc. Future experiments that scramble the sequence of the HZA rather than deleting it would better preserve topology and directly address this alternative model.

Deleting the HZA^mT^ enhancer had more complex effects on cadmium activated transcription. While it reduced activation of both adjacent genes *mtl-2* and *T08G5.1*, these genes still displayed residual activation in response to cadmium. These results indicate the existence of mechanisms to activate *mtl-2* and *T08G5.1* that function independently of the HZA^mT^ enhancer. This is consistent with the results of Earley et al (2021) showing that a subset of cadmium-activated genes do not require *hizr-1*. Furthermore, cadmium-activated genes positioned at a distance from this deletion displayed reduced activation in response to cadmium. We speculate that the enhancer deletion, by reducing the levels of *mtl-2* and/or *T08G5.1*, may influence cadmium activated transcription of genes throughout the genome. Another possible model is that the deletion strain contains a background mutation that affects cadmium-activated transcription. In this case, the background mutation does not appear to affect transcription regulated by excess zinc.

Deleting the HZA^cF^ enhancer partially reduced activation of both adjacent genes *cdr-1* and *F35E8.10* in response to cadmium. By contrast, control genes positioned at a distance from this deletion that are regulated by cadmium were minimally affected. These results support the conclusions that a single HZA enhancer can regulate two genes that are oriented head-to-head, and the HZA^cF^ enhancer can function in either orientation (Fig. 4E-F). As discussed above, this 14 bp deletion mutation also changed the spacing, and potentially the topology, of sequence elements in the promoter. Thus, an alternative explanation is that the deletion reduced cadmium-activated transcription as a result of this topology change. It was more difficult to interpret the results with excess zinc. Activation of *cdr-1* in response to excess zinc was significantly reduced by deleting the HZA^cF^ enhancer, demonstrating that the enhancer is necessary for zinc-activated transcription of the adjacent gene *cdr-1*. However, because *F35E8.10* transcripts in WT animals did not accumulate to a significant level in response to excess zinc, we were unable to evaluate the function of the HZA^cF^ enhancer in the regulation of *F35E8.10*. Our analysis using genome engineering to delete endogenous HZA enhancers supports and extends previous studies of HZA enhancers performed with extrachromosomal arrays (Roh et al. 2015; Earley et al. 2021).

Enhancers can function at a distance, but their effectiveness may be reduced by increasing distance (Quintero-Cadena and Sternberg 2016). We noticed a correlation between distance and magnitude of transcript accumulation in these two pairs of head-to-head genes. The *F35E8.10* start codon is 908bp from the HZA^cF^ enhancer, and the gene is weakly induced; the *mtl-2* and *T08G5.1* start codons are about 280bp from the HZA^mT^ enhancer, and these genes are induced at an intermediate level; the *cdr-1* start codon is 108bp from the HZA^cF^ enhancer, and the gene is strongly induced (Fig. 1D, H; Fig. S4A-B). These correlations do not rigorously establish a causal effect, and there may be reasons other than distance that dictate the magnitude of the response. A further caveat is that the two HZA enhancers are different, and they may have different relationships between distance and magnitude. Our results establish two experimental systems that can be used to directly investigate the relationship between distance to enhancer and magnitude of effect. For example, the HZA^cF^ enhancer could be engineered at different positions in the deletion mutant background to determine the impact on transcription of the head-to-head genes.

A single enhancer/suppressor can regulate multiple genes on the same strand, referred to as “enhancer sharing” (Fig. 4F top) (Quintero-Cadena and Sternberg 2016; Mills et al. 2020). However, little is understood about the ability of a single enhancer to control two genes that are oriented head-to-head. One factor that affects enhancer function is cell type. If the two head-to-head genes are expressed in different cell types, then the enhancer might exhibit different cell type-specific functions (Fig. 4F middle). Indeed, two groups reported orientation-dependent enhancer functions in mice and chordate *Ciona intestinalis* in different cell types (Swamynathan and Piatigorsky 2002; Hozumi et al. 2013). This cell type-specificity may be controlled by other nearby enhancer elements. In *C. elegans*, the HZA element is often in close proximity to a GATA element, an intestine-specific enhancer (Roh et al. 2015). At the *mtl-2*/*T08G5.1* locus, there are two predicted GATA elements flanking the HZA element (Fig. 4B) (Moilanen et al. 1999; Roh et al. 2015). Another factor that affects enhancer function is chromatin regulators, such as insulators that block access of RNA polymerase II to one flanking gene (Luan et al. 2022; Hamamoto et al. 2023). A lack of CTCF homologs in *C. elegans* may explain the ability of the HZA enhancer to function on both head-to-head genes (Heger et al. 2009). Our results indicate that the HZA^mT^ and HZA^cF^ enhancers can regulate two genes in the same conditions (Fig. 4F, bottom).

How does a single enhancer control two genes? A widely supported model for enhancer function is looping: proteins recruited to the enhancer and the promoter bring the two into a dynamic enhancer-promoter contact (EPC) (Panigrahi and O’Malley 2021). The dynamic EPC might be the rate-limiting factor in transcriptional activation of a target gene (Panigrahi et al. 2018). Thus, one possibility is that the HZA enhancer loops to one flanking gene at a time, forming only one EPC; in this scenario, the two gene promoters might compete for access to the EPC. To test this model, we deleted the *mtl-2* gene and examined transcription of *T08G5.1*.

Interestingly, transcription of *T08G5.1* was similar in the deletion mutant, indicating that the EPC may not alternate between the two promoters. Perhaps the EPC interacts with the promoters of both genes at the same time. Future experiments analyzing a deletion of *T08G5.1* would further test this relationship. MDT-15, a mediator subunit, is a co-activator for HIZR-1, and may contribute to EPC formation (Shomer et al. 2019).

There are several possible reasons why this orientation of enhancers and promoters may have evolved: 1) It is beneficial to have co-expression of two functionally related genes in response to metal stress. If two head-to-head genes function in the same pathway, then a single enhancer might be an efficient mechanism for mounting a response. 2) The “Sheltered Island Hypothesis” speculates that functionally unrelated genes remain clustered or nested together because any changes would be toxic to the essential genes in the cluster (Chen and Stein 2006). 3) One of the head-to-head genes may be a bystander gene that is neither beneficial nor toxic. If co-regulation of these genes is not deleterious, then this arrangement may have evolved. Whereas the *mtl-2* gene has been characterized extensively for a function in cadmium resistance, the function of the *T08G5.1* gene has not been characterized (Almutairi et al. 2024; Essig et al. 2024). In a similar manner, the *cdr-1* gene has been characterized for a function in cadmium resistance, but the function of the *F35E8.10* gene has not been characterized (Hall et al. 2012). Thus, all three models are possible explanations for this arrangement of genes.

## Data Availability

All strains and primers are available upon request. Supplementary File 1 contains all qPCR data processing spreadsheet (eg. Organized Data Cd, *hizr-1 am286*, Zn, etc), GraphPad Prism sheets, and log2 average calculation sheets (for superplot). Corresponding folders to figures: “*hizr-1 am286*” – Fig. 1, “Zn” – Fig. 2, Sup Fig. S1, S3, & S4, “Cd” – Fig. 3, Sup Fig. S2 & S4, “*mtl-2*”, “*srh-308*” – Fig. 4. Supplementary File 2 contains sequencing files for HZA^mT^ and HZA^cF^ loci. “*cdr-1*_sequencing.seq” and “*mtl-2*_sequencing.seq” are Sanger sequencing results for the knockout strains. “*mtl-2 gk125*” folder contains Sanger sequencing results for *gk125* allele verification. Supplementary File 3 contains sequences of *mtl-2* homologs from *C. breneri*, *C. briggsae*, and *C. remanei* used for alignment in Fig. 2. Files are made available online at https://zenodo.org/records/13948425, DOI: 10.5281/zenodo.13948425.

## Acknowledgement

We thank Barak Cohen and Mike Nonet for helpful discussions and the *Caenorhabditis* Genetics Center and SUNY Biotech for providing strains. Brian Egan, Dana Shaw, and Vishnu Saraswathy provided advice and assistance with qPCR procedures and data analysis.

