## Supplementary Tables and Figures for "A single high-zinc activation enhancer can control two genes orientated head-to-head in *C. elegans*"

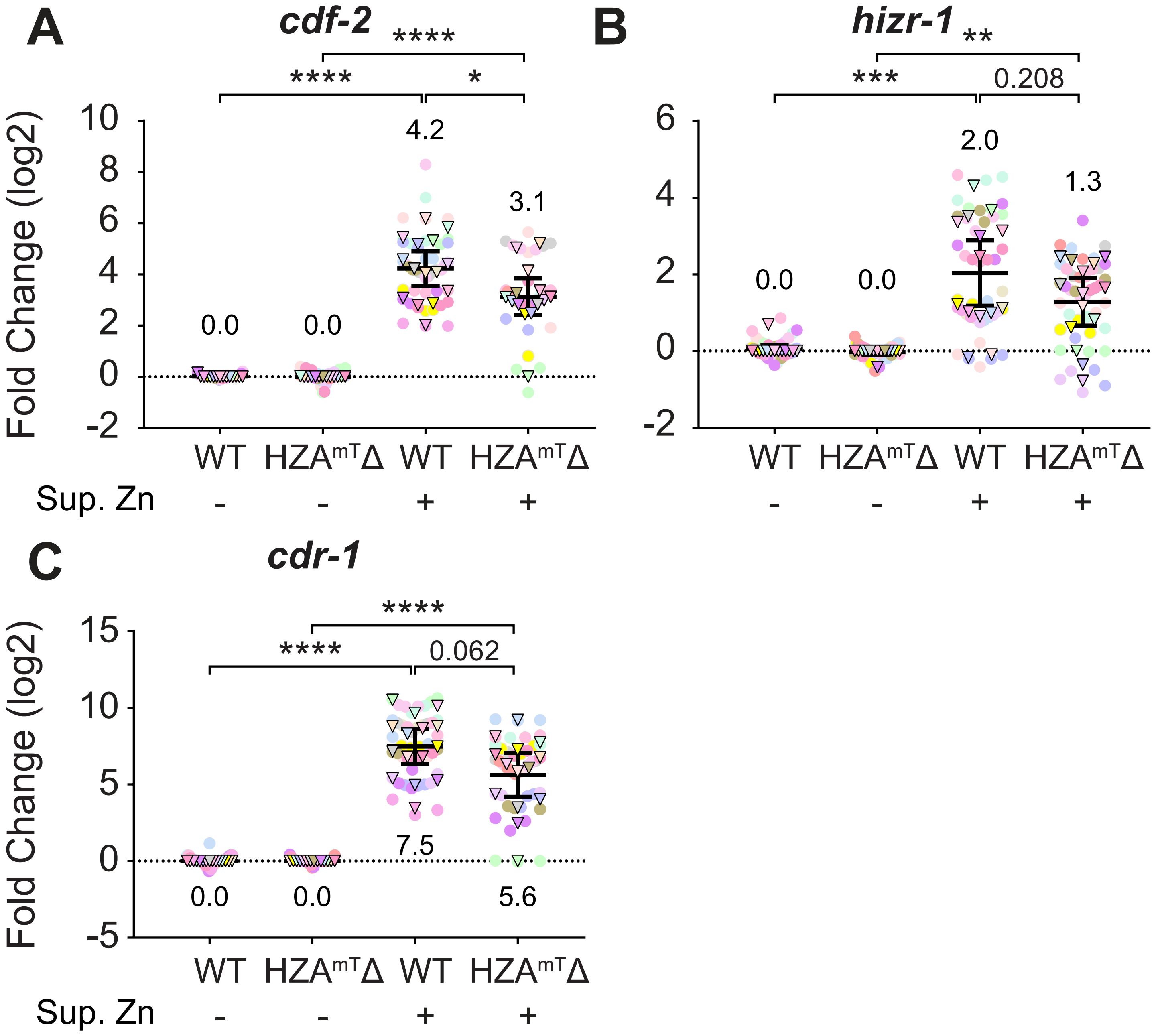

A
cdf-2
****
****
*
Fold Change (log2)
10
8
6
4
2
0
-2
4.2
3.1
0.0
0.0
WT
HZA$^{mT}$Δ
WT
HZA$^{mT}$Δ
Sup. Zn
-
-
+
+
B
hizr-1
**
***
0.208
6
4
2
0
-2
2.0
1.3
0.0
0.0
WT
HZA$^{mT}$Δ
WT
HZA$^{mT}$Δ
-
-
+
+
C
cdr-1
****
****
0.062
Fold Change (log2)
15
10
5
0
-5
7.5
0.0
0.0
5.6
WT
HZA$^{mT}$Δ
WT
HZA$^{mT}$Δ
Sup. Zn
-
-
+
+

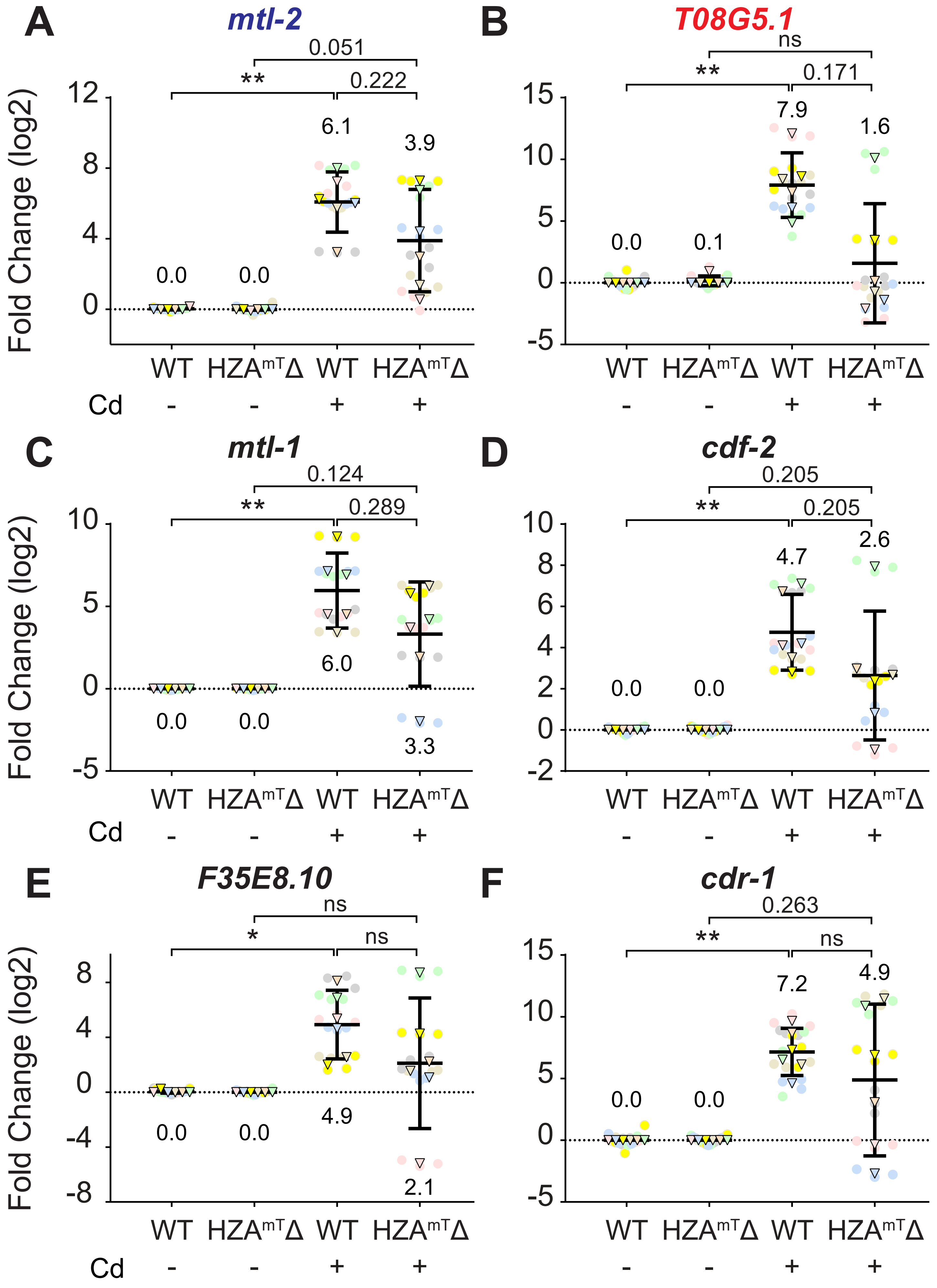

A
mtl-2
0.051
**
0.222
6.1
3.9
0.0
0.0
Fold Change (log2)
12
8
4
0
WT HZA^mT^Δ WT HZA^mT^Δ
Cd - - + +
B
T08G5.1
ns
**
0.171
7.9
1.6
0.0
0.1
15
10
5
0
-5
WT HZA^mT^Δ WT HZA^mT^Δ
- - + +
C
mtl-1
0.124
**
0.289
6.0
0.0
0.0
3.3
Fold Change (log2)
10
5
0
-5
WT HZA^mT^Δ WT HZA^mT^Δ
Cd - - + +
D
cdf-2
0.205
**
0.205
4.7
2.6
0.0
0.0
10
8
6
4
2
0
-2
WT HZA^mT^Δ WT HZA^mT^Δ
- - + +
E
F35E8.10
ns
*
ns
4.9
0.0
0.0
2.1
Fold Change (log2)
8
4
0
-4
-8
WT HZA^mT^Δ WT HZA^mT^Δ
Cd - - + +
F
cdr-1
0.263
**
ns
7.2
4.9
0.0
0.0
15
10
5
0
-5
WT HZA^mT^Δ WT HZA^mT^Δ
- - + +

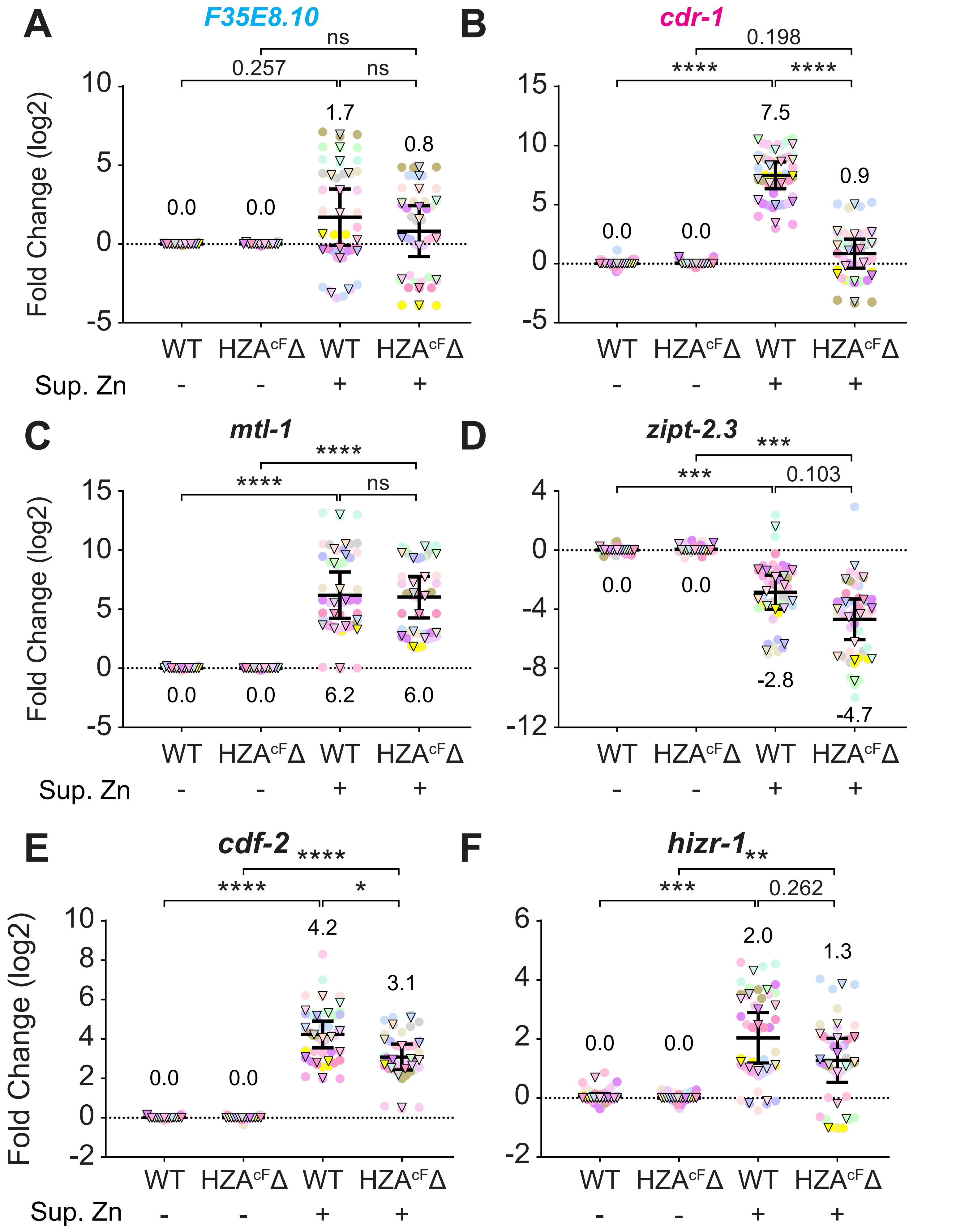
A
F35E8.10
ns
0.257
ns
1.7
0.8
0.0
0.0
Fold Change (log2)
WT
HZA$^{cF}$Δ
WT
HZA$^{cF}$Δ
Sup. Zn
-
-
+
+
B
cdr-1
0.198
****
****
7.5
0.9
0.0
0.0
C
mtl-1
****
****
ns
0.0
0.0
6.2
6.0
D
zipt-2.3
***
***
0.103
0.0
0.0
-2.8
-4.7
E
cdf-2
****
****
*
4.2
3.1
0.0
0.0
F
hizr-1
**
***
0.262
2.0
1.3
0.0
0.0

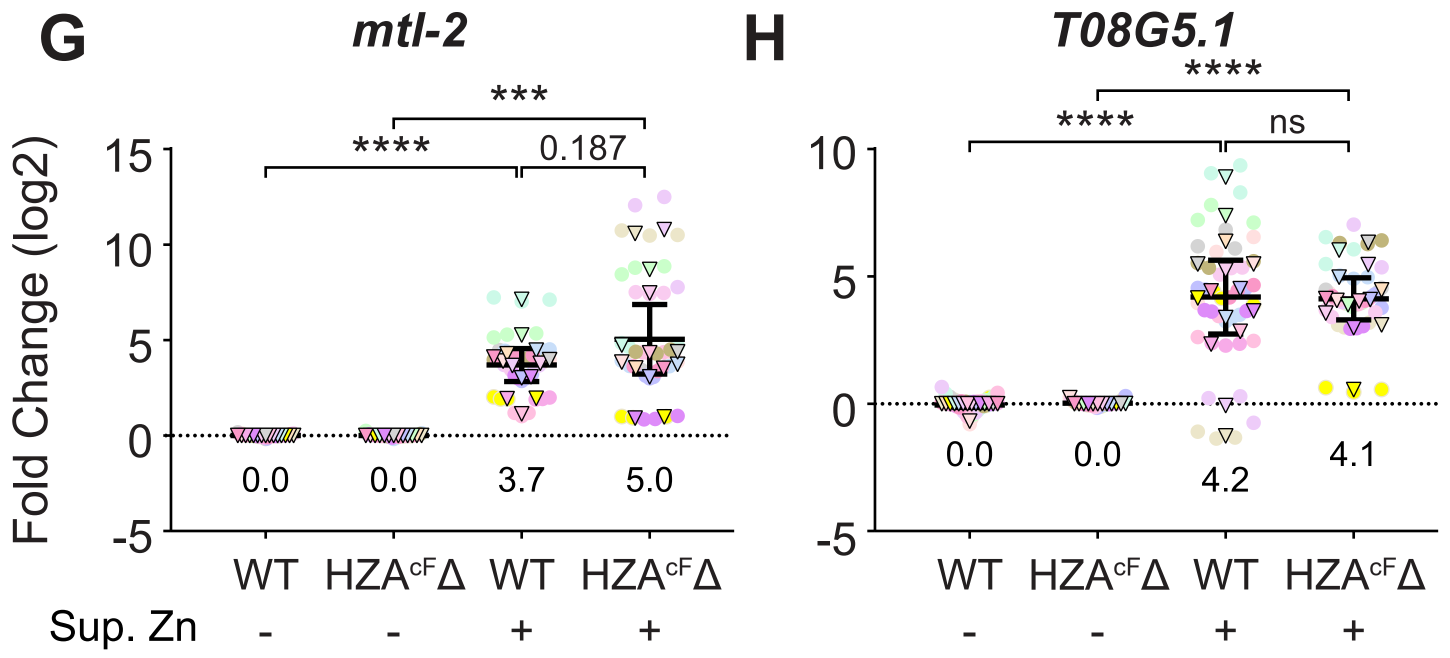
G
mtl-2
H
T08G5.1
Fold Change (log2)
15
10
5
0
-5
***
****
0.187
0.0
0.0
3.7
5.0
****
****
ns
0.0
0.0
4.2
4.1
WT
HZAcFΔ
WT
HZAcFΔ
Sup. Zn
-
-
+
+

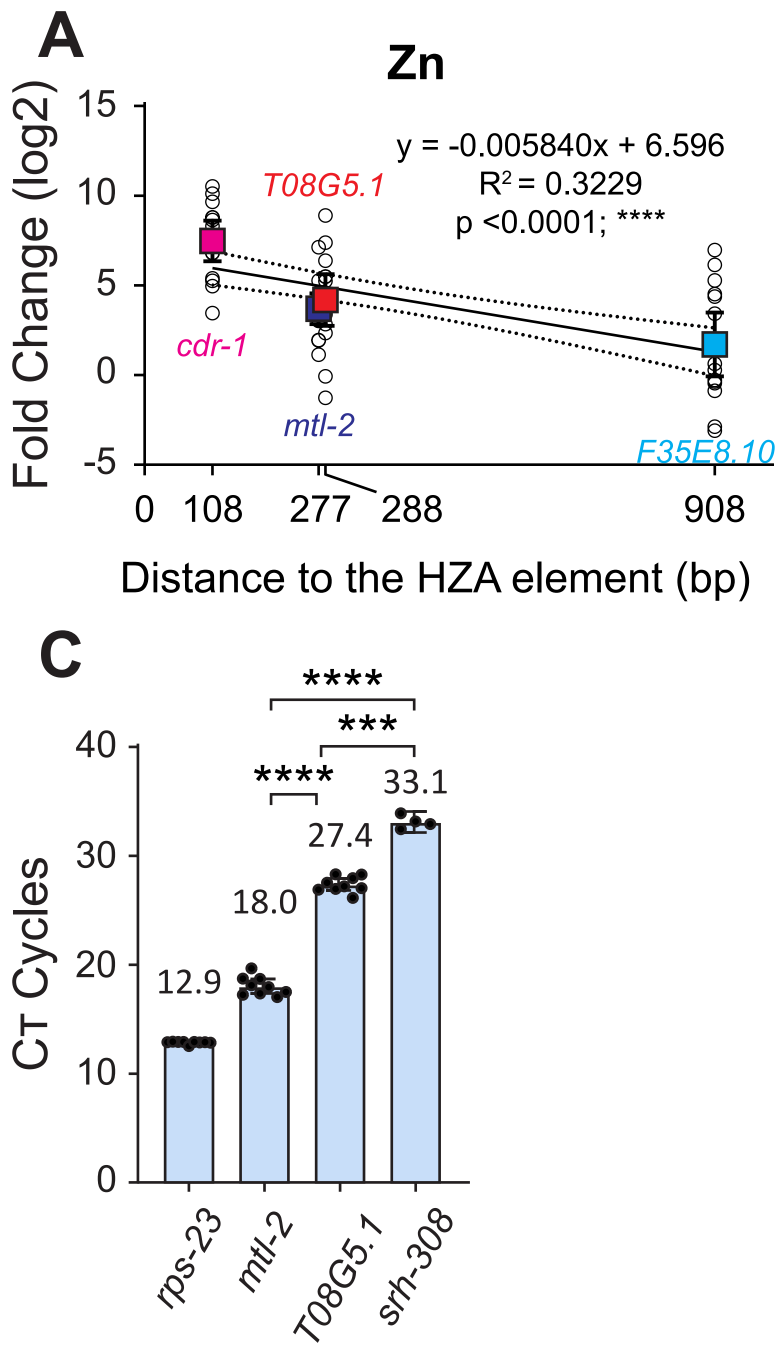
A
Zn
Fold Change (log2)
y = -0.005840x + 6.596
R² = 0.3229
p <0.0001; ****
T08G5.1
cdr-1
mtl-2
F35E8.10
15
10
5
0
-5
0
108
277
288
908
Distance to the HZA element (bp)
B
Cd
T08G5.1
y = -0.00295x + 7.68
R² = 0.162
p = 0.0511
cdr-1
mtl-2
F35E8.10
15
10
5
0
0
108
277
288
908
Distance to the HZA element (bp)
C
CT Cycles
****
***
****
40
30
20
10
0
12.9
18.0
27.4
33.1
rps-23
mtl-2
T08G5.1
srh-308

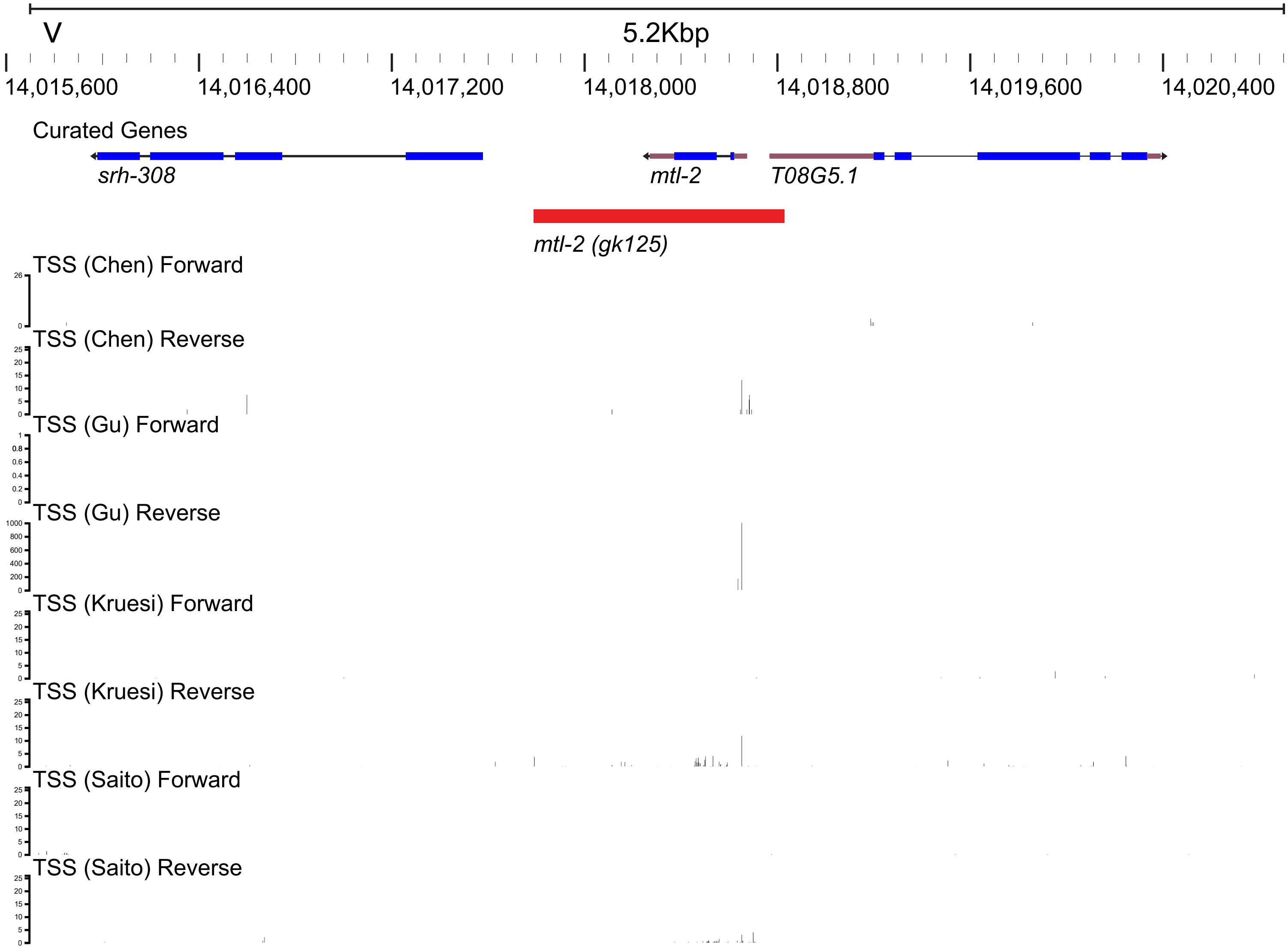
V
5.2Kbp
14,015,600
14,016,400
14,017,200
14,018,000
14,018,800
14,019,600
14,020,400
Curated Genes
srh-308
mtl-2
T08G5.1
mtl-2 (gk125)
TSS (Chen) Forward
TSS (Chen) Reverse
TSS (Gu) Forward
TSS (Gu) Reverse
TSS (Kruesi) Forward
TSS (Kruesi) Reverse
TSS (Saito) Forward
TSS (Saito) Reverse

**Supplementary Figure 1 (with main Figure 2).** The $HZA^{mT}$ element was not necessary for zinc-induced transcript accumulation of *cdf-2*, *hizr-1*, and *cdr-1*. **A-C)** Wild type (WT) and *syb4265* ($HZA^{mT}\Delta$) mutants at the L4 stage were cultured with replete (-) or 200µM supplemental zinc (+) for 16 hours and analyzed by qPCR. Values for WT and *syb4265* mutants with replete zinc were set equal to 0, and values with supplemental zinc represent log2 fold change. N = 15 initial biological replicates, but may vary in panels due to outlier removal. Circles are technical replicates, and triangles are biological replicates. Same color denotes the same experiment trial. Statistical analysis by pairwise one-way ANOVA. Non-significant p-values are listed for p<0.3; otherwise, "ns". For significant p-values: *<0.05; **<0.01; ***<0.001; ****<0.0001. Error bars represent mean ± 95% confidence intervals. Mean values are listed. All fold changes are in base 2 logarithm (e.g. 3 fold = 8x). All three genes were significantly induced by excess zinc in $HZA^{mT}\Delta$ mutants, but the level of mRNA accumulation in excess zinc was slightly lower in the mutant compared to WT. This difference was significant for *cdf-2* and a trend that was not significant with this sample size for *hizr-1* and *cdr-1*. We speculate that the $HZA^{mT}\Delta$ mutation indirectly affects the transcription of distant genes in excess zinc by reducing expression of *mtl-2*.

**Supplementary Figure 2 (with main Figure 2).** The $HZA^{mT}$ element was necessary for full cadmium-induced transcription of *mtl-2* and *T08G5.1*. **A-F)** Wild type (WT) and *syb4265* mutants at the L4 stage were cultured with or without 100µM cadmium for 16 hours and analyzed by qPCR. Values for WT and *syb4265* mutants with standard medium were set equal to 0, and values with cadmium represent log2 fold change. N = 6 initial biological replicates, but may vary in panels due to outlier removal. In WT animals *mtl-2* and *T08G5.1* displayed significant mRNA accumulation in response to cadmium. By contrast, in $HZA^{mT}\Delta$ mutants these genes did not display significant induction, indicating the $HZA^{mT}$ element is necessary for full cadmium-activated transcription. However, these genes did display a trend towards weak induction that was not significant with this sample size. In addition, the values for WT+ cadmium and $HZA^{mT}\Delta$ mutant +cadmium were not significantly different, consistent with the model that there is residual activation in response to cadmium. Similar results were observed with *mtl-1*, *cdf-2*, *F35E8.10*, and *cdr-1*. We speculate that the $HZA^{mT}\Delta$ mutation indirectly affects the transcription of distant genes during cadmium exposure by reducing the expression of *mtl-2*.

**Supplementary Figure 3 (with main Figure 3).** The HZA$^{cF}$ element was necessary for zinc-induced transcript accumulation of the adjacent gene *cdr-1* but not distantly positioned genes *mtl-1*, *zipt-2.3*, *cdf-2*, *hizr-1*, *mtl-2*, and *T08G5.1*. **A-H)** Wild type (WT) and *syb4134* mutants at the L4 stage were cultured with replete (-) or 200µM supplemental zinc (+) for 16 hours and analyzed by qPCR. Values for WT and *syb4134* mutants with replete zinc were set equal to 0, and values with supplemental zinc represent log2 fold change. N = 15 initial biological replicates, but may vary in panels due to outlier removal. The adjacent gene *cdr-1* was significantly induced by excess zinc, and the response was severely reduced by deleting the HZA$^{cF}$ element. Thus, the HZA$^{cF}$ element was necessary for this induction. The adjacent gene *F35E8.10* was not significantly induced by excess zinc in WT, so it is not possible to draw a conclusion about the necessity of the HZA$^{cF}$ element. Five distantly positioned genes were significantly induced by excess zinc in WT and *syb4134* mutants. For *mtl-1*, *hizr-1*, *mtl-2*, and *T08G5.1*, the level of induction was not significantly different in WT and *syb4134* mutants. For *cdf-2,* the level of induction was significantly lower in *syb4134* mutants compared to WT, indicating that the HZA$^{cF}$ enhancer may indirectly promote full activation of *cdf-2*. The *zipt-2.3* gene was significantly repressed by excess zinc in WT and *syb4134* mutants.

**Supplementary Figure 4 (with main Figures 2-4).** Comparison of gene expression in excess zinc and cadmium **A, B)** Fold change in mRNA levels is plotted against the distance from the start codon to the HZA element as shown in Figure 1 panels D and H. Data source: zinc is from Fig. 2 and S3, and cadmium is from Fig. 3 and S2. Statistical analysis by linear regression. Circles are biological replicates. Squares are the mean of all biological replicates. Error bars represent 95% confidence intervals. Black line is the regression line. Dotted line represents 95% confidence interval of the regression line. Induction by zinc displayed a significant p-value less than 0.0001, indicating an inverse correlation between distance and induction. However, the $R^2$ value of 0.3229 indicates a loose fit to the regression line. Induction by cadmium displayed a p-value of 0.0511, indicating a trend towards an inverse correlation between distance and induction that was not significant with this sample size. The $R^2$ value of 0.162 indicates the model does not fit the dataset well. A caveat to this analysis is that it combines data from two different HZAs. **C)** Bars represent $C_T$ values for indicated genes analyzed in WT animals cultured with standard medium for 16 hours after the L4 stage. Source of data: Figure 4. Dots represent biological replicates. N = 9 for *rps-23*, *mtl-2*, and *T08G5.1*; N = 4 for *srh-308*. Statistical analysis by Welch's t test. The lowest value was *rps-23*, an abundantly expressed gene used for normalization. The highest value was *srh-308*, indicating a low level of transcript accumulation.

**Supplementary Figure 5 (with main Figure 4)** Transcription start sites (TSS) of *srh-308*, *mtl-2*, and *T08G5.1* in relation to the *mtl-2(gk125)* deletion. Figure is adapted from wormbase JBrowse 2. We used the assembly c_elegans_PRJNA13758 as the reference genome and used its default curated gene track (abridged to show only *srh-308*, *mtl-2*, and *T08G5.1*). We loaded 4 tracks based on datasets in Gu et al. (2012), Chen et al. (2013), Kruesi et al. (2013), and Saito et al. (2013). (under "Misc"->"Transcription Start Sites"). We also loaded change-of-function alleles (under "Alleles, Variations, RNAi"). At the top is a black line indicating a 5.2 kbp region on Chromosome V, and below that are the coordinates. Each long tick is indicated by its basepair position, and the interval between two long ticks is 800bp. The dataset tracks are listed and labeled below. Curated genes show the position and the orientation of each gene (black arrowheads), as well as 5' UTR (grey boxes), exons (blue boxes), and introns (black lines). Below that is an abridged track showing the *gk125* deletion (red bar). Each TSS dataset contains 2 tracks: the forward strand and the reverse strand. Each identified TSS position is indicated by a bar, the height of which is its confidence score in linear scale. The scale bars of the scores are on the left of each track. TSS (Gu) Forward track does not contain any TSS position calls in this region. *gk125* removes most of the identified TSS positions in this region, including high-confidence consensus positions (presumed to be for *mtl-2*) and multiple lower-confidence non-consensus positions (presumed to be for *srh-308*). The deletion of *srh-308* TSS might explain why *srh-308* was not induced by excess zinc in *mtl-2(gk125)* mutant animals.

Online sources for this figure:

**Wormbase JBrowse**: https://wormbase.org/tools/genome/jbrowse-simple/?data=data%2Fc_elegans_PRJNA13758&loc=V%3A14015787..14021917&tracks=Curated_Genes%2CTSS%20(Gu)%20Forward%2CTSS%20(Gu)%20Reverse%2CTSS%20(Chen)%2

0Forward%2CTSS%20(Chen)%20Reverse%2CTSS%20(Saito)%20Forward%2CTSS%20(Saito)%20Reverse%2CTSS%20(Kruesi)%20Forward%2CTSS%20(Kruesi)%20Reverse%2CChange-of-function%20alleles&highlight=

**Wormbase JBrowse 2**: https://wormbase.org/tools/genome/jbrowse2/?session=share-Q7V4jUzqIT&password=Y1H5S

**Supplementary Table 1: *C. elegans* Strain List**

| **Strain Name**[1] | **Genotype**[2] | **Comments** |
|---|---|---|
| N2 | Wild type | Bristol isolate (Brenner 1974) |
| WU946 | *mtl-2(gk125)* V | Deletion of a region from 208bp upstream of *mtl-2* ATG start codon to 584bp downstream of *mtl-2* stop codon; insertion of one adenine; outcrossed 5 times to N2 from VC128 (The *C. elegans* Deletion Mutation Consortium, 2012) |
| WU1958 | *hizr-1(am286)* X | Q87 to STOP codon (C259T) in *hizr-1* (Warnhoff et al. 2017) |
| PHX4134 (WU1993) | *syb4134* V | CRISPR-generated deletion of the $HZA^{cF}$ element, AAC AGA AAC TAC AA, positioned 108bp upstream of the *cdr-1* start codon and 908bp upstream of the *F35E8.10* start codon. Confirmed by DNA sequencing (primers listed in Table S2). Generated by SunyBiotech (www.sunybiotech.com) |
| PHX4265 (WU1992) | *syb4265* V | CRISPR-generated deletion of the $HZA^{mT}$ element, ATC ACA AAC TAG AGT, 278bp upstream of *mtl-2* start codon and 287bp upstream of *T08G5.1* start codon. Confirmed by DNA sequencing (primers listed in Table S2). Generated by SunyBiotech (www.sunybiotech.com) |

[1] Strain names are standard *C. elegans* designations that describe the complete genotype in a single name.
[2] The alleles of interest in the strain and their chromosomal location.

**Supplementary Table 2: Oligonucleotide Primers for qPCR and DNA sequencing**

| Gene[1] | Forward (5'-3')[2] | Reverse (5'-3')[2] | Product size (bp)[3] |
|---|---|---|---|
| *ama-1* qPCR | ATC GGA GCA GCC AGG AAC TT | GAC TGT ATG ATG GTG AAG CTG G | 98 |
| *cdf-2* qPCR | CAA GAT TGC ACT CTC CGT ACA | TTT CGA GTT TCG CGT AGA ATT G | 75 |
| *cdr-1* qPCR | GAC TCA AGA ACT GTC CGA ACT T | CGT TCA GAG GTG CAT GTG ATA | 103 |
| *F35G8.10* qPCR | CTG GTT ACA ACT GCC CAA ATG | GAC AGT CCG CAG CGA TAA T | 108 |
| *hizr-1* qPCR | TCA TTT TGC GGT TTC ATC GTG | CAT CGC GTG TAT CTA CAG CTA C | 150 |
| *mtl-1* qPCR | TGG ATG TAA GGG AGA CTG CAA | CAT TTT AAT GAG CCG CAG CA | 107 |
| *mtl-2* qPCR | CCA AGT GCT GTG AGC AAT ACT | CCT GAG CAC ATT CGC AGT T | 105 |
| *rps-23* qPCR | CGT GTC CAG CTC ATC AAG AA | AAC GTC CGA AAC CAG ATA CG | 102 |
| *T08G5.1* qPCR | CAT TAT GTG GCC CAG GAA CT | CCT GAA GAC GAG CTT CAC TAA A | 107 |
| *zipt-2.3* qPCR | TGG ATT ATG CTC GGA GTT ATG G | CGT CGT GTT CAG GTA GGA AAT | 84 |
| *syb4134* DNA sequencing | TGC GGA AAG AAC AAT TGA AA | AGT AAA TGA CAG CGG CTC CA | 391 in N2 |
| *syb4265* DNA sequencing | TCA GTG GAA CGA AAC AAA CG | GCG TGC ACA ATT ATT TGG ATT | 357 in N2 |
| *gk125* DNA sequencing | TCA GTG GAA CGA AAC AAA CG | GCA GTT GGG CAG CAG TAT | 719 in N2 |
| *mtl-2* genotyping | ATG TTC TGG AGC GGT TCT GG | GAA CGG TCA CTT CGA TGG CT | 1747 in N2 |

[1] The target gene or allele for the primers
[2] The forward and reverse primers used to amplify the region around or within the genes or mutant alleles
[3] The size of the fragment from PCR using the primers listed with wild type (N2)
